# Intermediate filaments control collective migration by restricting traction forces and sustaining cell-cell contacts

**DOI:** 10.1101/328609

**Authors:** Chiara De Pascalis, Carlos Pérez-González, Shailaja Seetharaman, Batiste Boëda, Benoit Vianay, Mithila Burute, Cécile Leduc, Nicolas Borghi, Xavier Trepat, Sandrine Etienne-Manneville

**Affiliations:** Institut Pasteur Paris CNRS UMR3691, Cell Polarity, Migration and Cancer Unit, INSERM, Equipe Labellisée Ligue Contre le Cancer, 25 rue du Dr Roux 75724, Paris Cedex 15, France.; Sorbonne Universités, UPMC Univ Paris 06, IFD, 4 Place Jussieu, 75252 Paris cedex 05, France; Institute for Bioengineering of Catalonia, Barcelona Institute of Science and Technology (BIST), Barcelona 08028, Spain.; Facultat de Medicina, University of Barcelona, Barcelona 08028, Spain; Université Paris Descartes, Sorbonne Paris Cité, Paris, 75006, France.; University of Paris Diderot, INSERM, CEA, Hôpital Saint Louis, Institut Universitaire d’Hematologie, UMRS1160, CytoMorpho Lab, 75010, Paris, France and University of Grenoble-Alpes, CEA, CNRS, INRA, Biosciences & Biotechnology Institute of Grenoble, Laboratoire de Phyisologie Cellulaire & Végétale, CytoMorpho Lab, 38054, Grenoble, France; Institut Jacques Monod, Unité Mixe de Recherche 7592, Centre National de la Recherche Scientifique, Université Paris-Diderot, Paris 75013, France; Institució Catalana de Recerca i Estudis Avançats (ICREA), Barcelona, Spain.; Centro de Investigación Biomédica en Red en Bioingeniería, Biomateriales y Nanomedicina, 08028, Spain; CytoMorpho Lab^6^

## Abstract

Mesenchymal cell migration relies on the coordinated regulation of the actin and microtubule networks which participate in polarised cell protrusion, adhesion and contraction. During collective migration, most of the traction forces are generated by the acto-myosin network linked to focal adhesions at the front of leader cells, which transmit these pulling forces to the followers. Here, using an *in vitro* wound healing assay to induce polarisation and collective directed migration of primary astrocytes, we show that the intermediate filament (IF) network composed of vimentin, GFAP and nestin contributes to directed collective movement by controlling the distribution of forces in the migrating cell monolayer. Together with the cytoskeletal linker plectin, these IFs control the organisation and dynamics of the acto-myosin network, promoting the actin-driven treadmilling of adherens junctions, thereby facilitating the polarisation of leader cells. Independently of their effect on adherens junctions, IFs influence the dynamics and localisation of focal adhesions and limit their mechanical coupling to the acto-myosin network. We thus conclude that IFs promote collective directed migration by restricting the generation of traction forces to the front of leader cells, preventing aberrant tractions in the followers and by contributing to the maintenance of lateral cell-cell interactions.

## Introduction

During morphogenesis, tissue repair and cancer, cells frequently migrate in a collective manner as groups, chains or sheets (Haeger et al., 2015; Mayor and Etienne-Manneville, 2016). Collectively migrating cells move with a similar speed and direction (Etienne-Manneville, 2014). The cytoskeleton, composed of actin microfilaments, microtubules and intermediate filaments (IFs), plays a key role in single and collective cell migration. The roles of the actin and microtubule network in cell migration have been well characterised (Etienne-Manneville, 2013; Gardel et al., 2010). Similar to actin and microtubules, IFs affect cell migration (Leduc and Etienne-Manneville, 2015). During oncogenesis, changes in IF protein expression, in particular increased vimentin levels, have been associated with cell invasion and tumour spreading (Chung et al., 2013; Leduc and Etienne-Manneville, 2015). However, the exact functions of IFs during migration are still not well understood.

Migration of cell sheets is characterised by specific mechanical features (De Pascalis and Etienne-Manneville, 2017). High tractions generated at the front of leader cells (du Roure et al., 2005; Trepat et al., 2009) are transmitted to the rest of the monolayer (Serra-Picamal, 2012; Tambe et al., 2011). Due to their structural characteristics, IFs are hypothesised to be key players in cell mechanics, helping maintain cell and tissue integrity (Herrmann et al., 2009; Kreplak and Fudge, 2007). Keratin IFs have recently been shown to control traction forces during collective migration of epithelial cells (Sonavane et al., 2017). Moreover, alterations of the vimentin network in endothelial cells perturb acto-myosin contractility and cell mechanical resilience (Osmanagic-Myers et al., 2015), suggesting that non-keratin IFs may also contribute to the mechanical properties of collectively migrating cells.

In this study, we use primary astrocytes to investigate the role of IFs in collective migration. Astrocytes are major glial cells that mainly express the three IF proteins Glial Fibrillary Acidic Protein (GFAP), nestin and vimentin, that assemble together in cytoplasmic IFs (Leduc and Etienne-Manneville, 2017). Increased levels of these IF proteins have been reported in glioblastomas, which are highly invasive primary glial tumours (Lin et al., 2016; Lv et al., 2017; Ma et al., 2008; Matsuda et al., 2015). Astrocytes migrate in a collective manner during development (Gnanaguru et al., 2013; Liu et al., 2015). In the adult brain, reactive astrocytes, which express higher levels of GFAP, can polarise to eventually migrate in the direction of inflammatory sites (Burda et al., 2016; Pekny et al., 2016). The collective migration of reactive astrocytes can be recapitulated *in vitro* in a wound healing assay which induces the polarisation of wound edge cells and the collective movement of the cell sheet in a timely controlled manner (Etienne-Manneville, 2006; Etienne-Manneville and Hall, 2001). Using this assay, we have investigated the role of vimentin, GFAP and nestin in collective migration. We show that these three IF proteins participate in the dynamics of the acto-myosin network and its association with focal adhesions (FAs) and adherens junctions (AJs). Glial IFs thus control the distribution of forces in the migrating monolayer and the interactions between neighbouring cells, ultimately determining the speed and direction of collective migration.

## Results and discussion

### IFs control the distribution and strength of traction forces in a migrating monolayer

We used previously described siRNAs (si triple IF) to specifically decrease the expression of each of the three main glial cytoplasmic IF proteins (GFAP, vimentin and nestin) expressed in primary rat astrocytes (Dupin et al., 2011). A second independent set of siRNAs was also used to reduce the amount of IFs in astrocytes (Fig. S1A, S1B). Upon IF protein depletion, cell area and size during migration was increased, as previously reported (Middeldorp and Hol, 2011)(Fig. S1C, S1D). In wound healing assays, IF-depleted cells (si triple IF) were slower than control cells (Fig. 1A, 1C, S1E and movie 1) confirming that IFs are involved in astrocyte migration as previously observed *in vitro* and *in vivo* (Dupin et al., 2011; Lepekhin et al., 2001; Sakamoto et al., 2013). Reduction of IFs also significantly decreased the persistence and directionality of migration of primary astrocytes (Fig 1B, 1C, S1E) and centrosome reorientation, an indicator of the front-rear polarisation of the leader cells (Fig. S1F). Depletion of either of the three IF proteins present in astrocytes had similar although less pronounced effects on migration than the concomitant depletion of the three proteins, suggesting that they all participate in the functions of the glial IF network (Fig. 1C).

**FIGURE 1:**
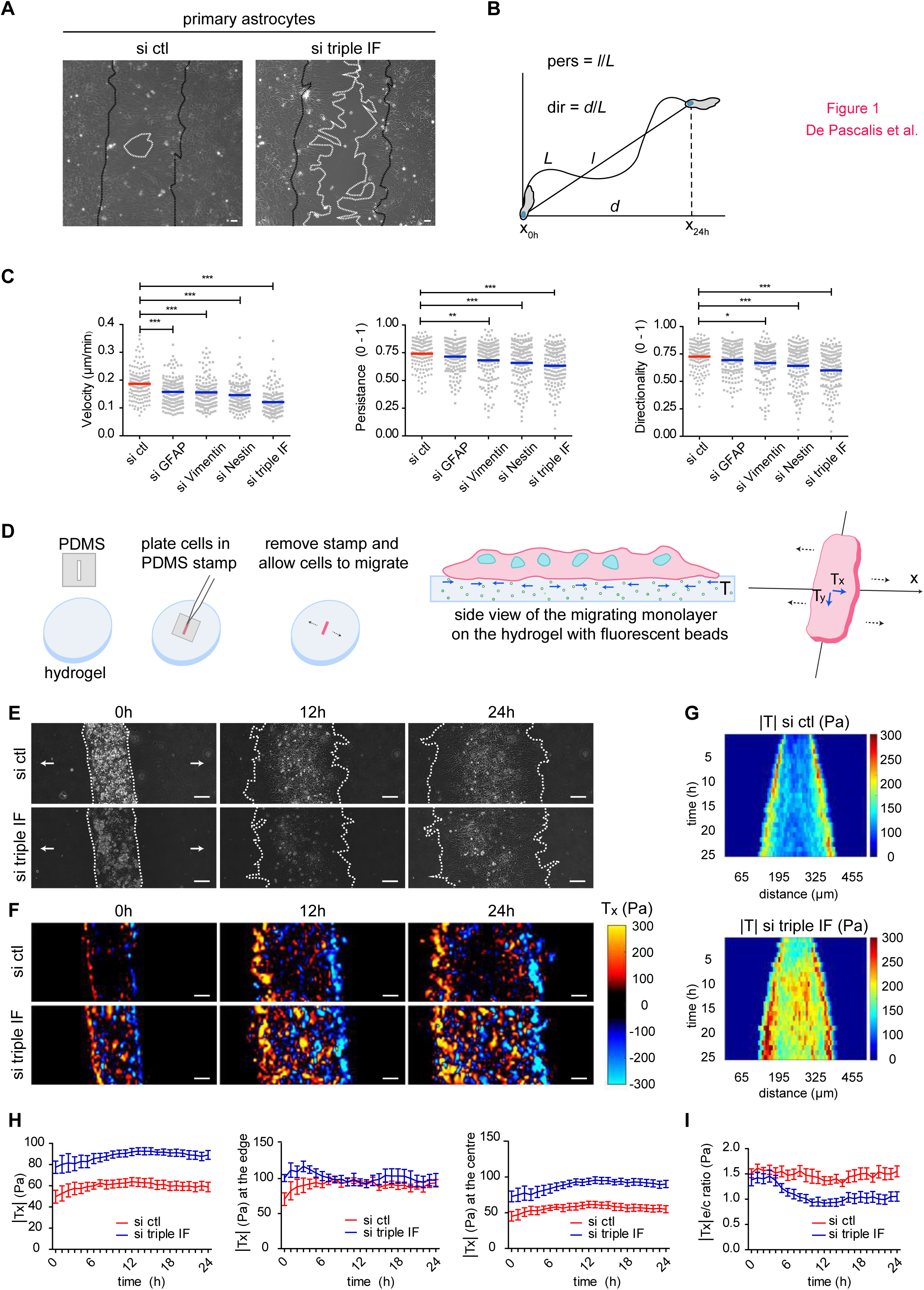
Intermediate filaments control collective astrocyte migration by regulating traction forces. **A)** Phase contrast images of astrocyte (shown in **Movie 1**) wound healing after 24h of migration. Black lines represent the initial position (0h) and white lines show the final position (24h) of the leading edge. **B)** Simplified method for calculating persistence and directionality of migration of a cell with nuclear tracking. For more detailed formulas, see Methods section. **C)** Graphs of cell velocity, directionality and persistence measured by manual nuclear tracking of leader cells after 24h of migration. **D)** Simplified protocol for plating cells into PDMS rectangular stamps onto hydrogels. The black dotted arrows show the main directions of migration. The central image shows a monolayer of cells migrating on a hydrogel embedded with fluorescent beads (green dots) with representative tractions (T, blue arrows). The last image represents schematically the components of tractions (T_x_ and T_y_) analysed with TFM. **E)** Phase contrast images of astrocytes migrating on a 9kPa collagen-coated polyacrylamide hydrogel at different time points. The white dotted line represents the leading edge, while the arrows show the direction of migration. See also **Movie 2**. **F)** Tractions in the x direction (T_x_) at indicated time points. See also **Movie 3**. **G)** Representative kymographs of total tractions (|T|). **H)** Graphs of tractions in the x direction (T_x_), average values (left), values at the edge of the monolayer (middle) and at the centre (right). **I)** Graph showing the ratio of the tractions at the edge over tractions at the centre, plotted as a function of time of migration. Data is from N = 3 independent experiments. The sample size for each repeat is: si ctl 50, 50, 50, si triple IF 50, 50, 50, si GFAP 50, 50, 50, si Vimentin 50, 50, 49, si Nestin 50, 50, 50 cells for 1C; 2, 1 and 4 videos of si ctl and 2, 3, 3 videos of si triple IF for 1E-I. Graphs show mean ± s.e.m. P values are * for p < 0.05, **: p for p < 0.01 and *** for p < 0.001. Scale bar 50 μm.

To analyse the effect of IF depletion on traction forces in a migrating astrocyte monolayer, we performed traction force microscopy (TFM) (Trepat et al., 2009) (Fig. 1D-1F and movie 2). Cells were plated on collagen-coated polyacrylamide substrates inside a polymethylsiloxane (PDMS) stamp of rectangular shape. Removal of the PDMS stamp allows the cells to migrate outwards (Fig. 1D, 1E and movie 2). Analysis of traction forces in control cells showed that most traction forces were exerted at the edges of the monolayer, whereas follower cells produced weaker forces (Fig. 1F-I and movie 3). Although in these conditions cell migration speed was not significantly reduced by IF depletion, possibly indicating that IF effects on velocity is sensitive to substrate rigidity, si triple IF cells showed stronger tractions at all time points (Fig. 1F-I). Moreover, in IF-depleted astrocytes, tractions were generated not only by leaders but also by followers. In fact, most of the increase in traction forces corresponded to increased forces in the follower cells resulting in the generation of similar traction forces at the edge and in the centre of the monolayer (Fig. 1F-I). These results show that IFs restrict the generation of traction forces to the leader cells and prevent the accumulation of traction forces within the monolayer.

### IFs control the organisation and dynamics of the acto-myosin network

In single cells, IFs have been shown to influence the organisation of the actin cytoskeleton as well as acto-myosin contractility (Costigliola et al., 2017; Gregor et al., 2014; Jiu et al., 2015; Jiu et al., 2017), leading us to investigate how IFs influence the acto-myosin network in migrating monolayers. During collective migration, astrocytes showed two main types of actin fibres (Fig. 2A and S2A). In addition to the longitudinal stress fibres oriented in the direction of migration and associated with FAs at the cell front, collectively migrating astrocytes display interjunctional transverse arcs (ITAs), which are anchored at AJs on the lateral sides of adjacent cells (Peglion et al., 2014). IF-depleted leader cells often completely lacked visible ITAs and only showed longitudinal stress fibres (Fig. 2A, 2B and S2A, S2B). Silencing any of the three IF proteins similarly favoured the generation of longitudinal stress fibres at the expense of ITAs (Fig. S2C). The general orientation of the actin cables was also affected in follower cells just behind the leaders (Fig. 2C, S2D). While control cells showed actin fibres both perpendicular and parallel to the wound, si triple IF cells mainly displayed stress fibres oriented perpendicularly to the wound both in leaders and followers (Fig. 2C, S2D).

**FIGURE 2:**
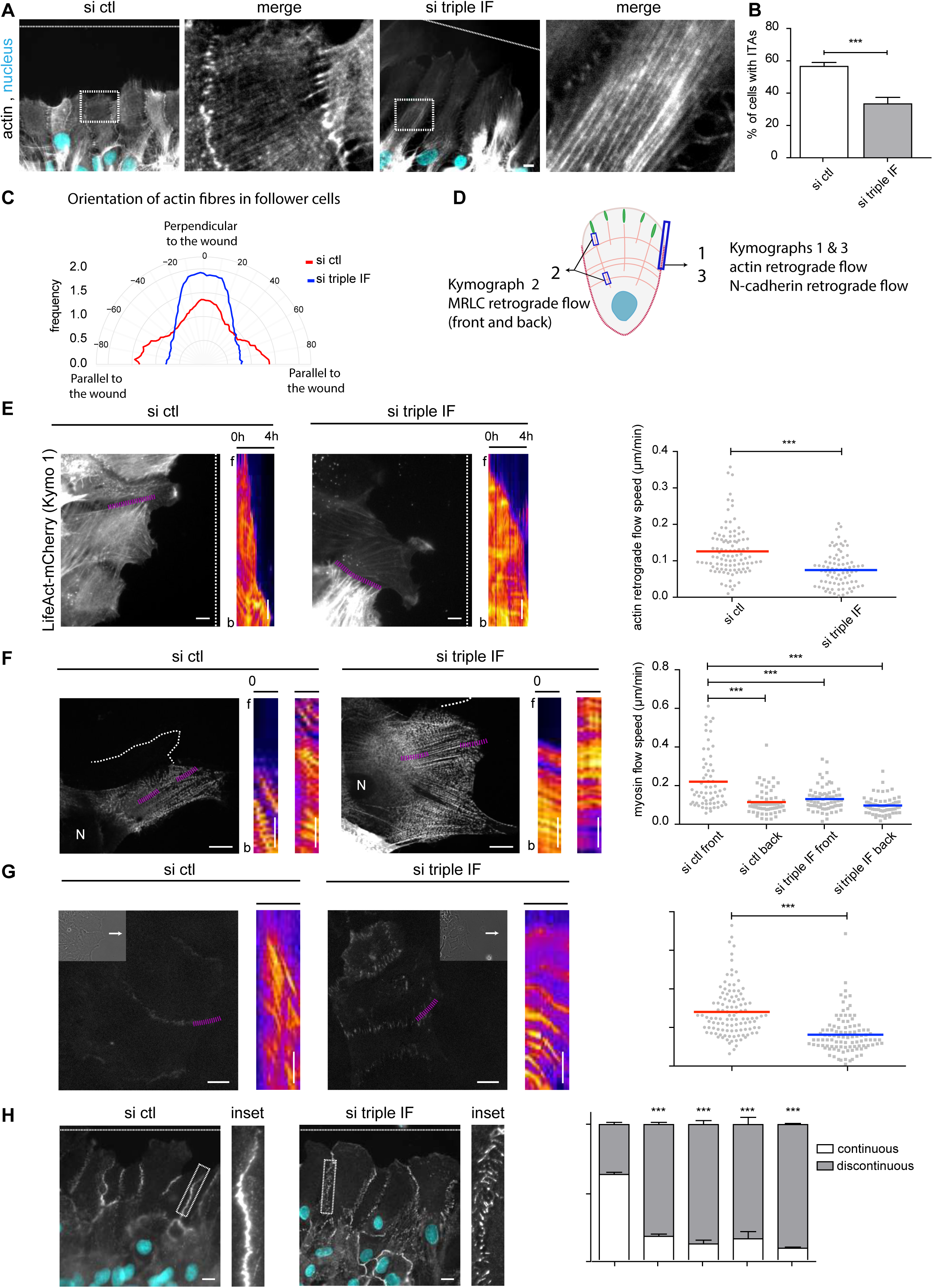
Intermediate filaments control the organisation and dynamics of the actin network. **A)** Immunofluorescence images of actin fibres in migrating astrocytes stained for nuclei (cyan) and actin (phalloidin, grey). **B)** Graph showing the percentage of leader cells that present interjunctional transverse arcs (ITAs) in si control (si ctl) and IF-depleted cells (si triple IF). **C)** Rose plot showing the frequency of angle distribution of actin fibres analysed in follower cells (Fig. S3D). Fibres with a 0° angle are perpendicular to the wound while fibres with an angle close to 90° are parallel to the wound. **D)** Schematics showing the position of the kymographs acquired in **E-G**. **E)** Frames of migrating si ctl and si triple IF astrocytes transfected with LifeAct-mCherry. The white dotted line represents the wound while the pink dotted lines represent the positions in which the kymographs were calculated. Kymographs (fire LUT) show the retrograde flow of actin on cell-cell junctions over a time period of 4h. The graph shows the mean retrograde flow speed of actin cables measured at the level of the cell-cell junctions. **F)** Frames from **Movie 4** of migrating astrocytes transfected with GFP-MRLC. The thick white dotted line represents the outline of nearby cells. The kymographs were obtained along the pink dotted lines. f and b on the side of the kymograph indicate the front and the rear of the line. Kymographs (fire LUT) show the retrograde flow of myosin at the front and at the rear of myosin longitudinal fibres over a time period of 15 min. The graph shows the mean speed of the myosin retrograde flow at the cell front and at the rear of the longitudinal fibre calculated from the kymographs. The white dotted lines indicate the position of the wound. **G)** Immunofluorescence images from **Movie 5** showing GFP-N-cadherin expressing astrocytes. Insets show the corresponding phase-contrast image. ‘N’ indicates the nucleus. The kymographs were obtained along the pink dotted lines. f and b on the side of the kymographs indicate the front and the rear of the line. Kymographs (fire LUT) show the retrograde flow of N-cadherin over a time period of 2h. The graph shows the mean retrograde flow speed of the N-cadherin flow in si ctl and si triple IF migrating astrocytes. **H)** Staining for N-cadherin (grey) and nuclei (cyan) in migrating astrocytes nucleofected with the indicated siRNAs. Histogram shows the mean percentage ±s.e.m of continuous junctions between adjacent leader cells. White dotted line indicates the position of the wound. Data is from N = 3 independent experiments. The sample size for each repeat is: si ctl 64, 67, 92 and si triple IF 44, 96, 122 for 2B; si ctl 11, 10, 8 stacks and si triple IF 12, 10, 8 stacks for 2C; si ctl 30, 40, 32 and si triple IF 18, 33, 30 for 2E; si triple IF f 22, 27, 18 and si triple IF b 22, 24, 20 for 2F; si ctl 14, 38, 28 and si triple IF 38, 49, 14 for 2G; si ctl 104, 120, 178 si GFAP 134, 137, 125, si vimentin 113, 180, 133, si nestin 132, 190, 123 and si triple IF 122, 225, 112 for 2H. P values are *** for p < 0.00. Scale bar 10 μm for all images except kymographs for which scale bar is 5 μm.

The dynamics of the acto-myosin network is one of the main factors responsible for the generation of traction forces exerted on the substrate. We used LifeAct-mCherry expressing astrocytes to analyse the dynamics of the actin network in leader cells. In wound edge cells, actin ITAs undergo a continuous retrograde flow coupled to the retrograde flow of associated AJs (Peglion et al., 2014). The retrograde flow of ITAs was visible in si ctl cells (Fig 2D, 2E). In the few si triple IF cells showing ITAs, the retrograde flow was much slower (Fig 2E). To analyse the retrograde flow in the longitudinal stress fibres, we used GFP labelled Myosin Regulatory Light Chain (GFP-MRLC) expressing cells (Fig 2D, 2F). Kymograph analysis at the front and rear end of these longitudinal fibres showed that retrograde flow of myosin was slower in IF-depleted cells than in control cells (Fig. 2D, 2F and Movie 4). Moreover, while myosin flow was higher at the front than at the rear of the fibres in control cells, such difference was abolished in si triple IF cells (Fig. 2F). Altogether, these results show that IFs play a key role in the organisation and dynamics of the acto-myosin network during collective cell migration.

### IFs are required for actin-driven treadmilling of AJs and maintenance of cell-cell interactions

The coordination of the actin-driven retrograde flow between adjacent leader cells drives the treadmilling of cadherin-based AJs which participates in the maintenance of AJs and thereby controls centrosome orientation and direction of migration in leader cells (Camand et al., 2012; Dupin et al., 2011; Peglion et al., 2014). Using GFP-N-cadherin expressing cells, we observed that depletion of IFs strongly reduced AJ retrograde flow, which was sometimes absent or in the opposite direction in some cells (Fig. 2D, 2G and movie 5). As previously described (Peglion et al., 2014), the reduction of AJ retrograde flow by si triple IF or by single siRNAs was associated with the altered morphology of AJs which appeared more discontinuous than in control cells (Fig. 2H, S2E, S2F). The alteration of the lateral AJs was confirmed by the increase of non-adherent regions between neighbouring wound-edge cells (Fig. S2F). Altogether, these results show that IFs facilitate the actin-driven dynamics of AJs between adjacent leader cells. The reduction of AJ retrograde flow in IF depleted cells along with the strong decrease of ITAs cables probably contributes to the decrease of cell polarity and to the loss of coordination between leader cells.

### IFs and plectin interact with FAs and control their dynamics and distribution

The impact of IFs on longitudinal stress fibres anchored at FAs (Fig. S2A) led us to investigate the relationship between IFs and FAs in migrating astrocytes expressing GFP-vimentin and paxillin-Orange (or Cherry-vimentin and GFP-paxillin). Live super-resolution (3D-Structured Illumination Microscropy, SIM) imaging showed that in astrocytes, like in other cell types (Burgstaller et al., 2010; Gonzales et al., 2001; Lynch et al., 2013; Mendez et al., 2010), IFs come in very close proximity to FAs (Fig. 3A). It also illustrated the highly dynamic behaviour of IFs near FAs (movies 6 and 7).

**FIGURE 3:**
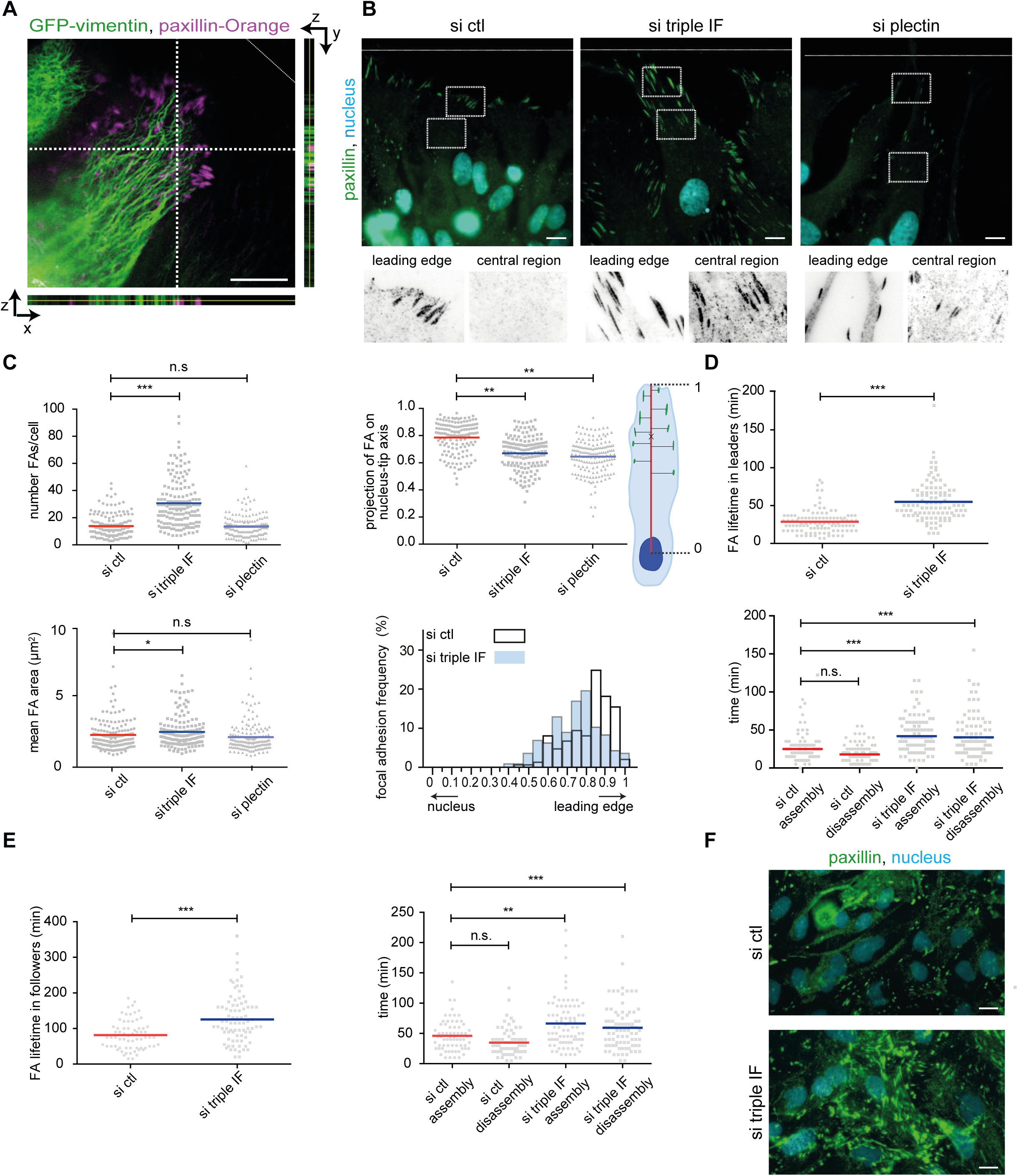
Intermediate filaments and plectin regulate focal adhesion localisation and dynamics. **A)** Fluorescence SIM-3D images of a migrating astrocytes transfected with GFP-vimentin and paxillin-Orange shown in **Movie 6** (see also **Movie 7**). The orthogonal projections show that vimentin and paxillin are found in the same focal plane. **B)** Immunofluorescence images of migrating astrocytes stained for nuclei (cyan) and paxillin (green). The white dotted lines indicate the position of the wound. Insets show enlarged images of focal adhesions at the leading edge and in a central region of the cell. **C)** The top left graph shows the mean number of focal adhesions per cell and the bottom left graph show the mean area of focal adhesions. Adhesions were projected on the nucleus-tip axis (see schematics). The central top graph shows the normalised distance to the nucleus centre of each FA. The central bottom graph shows the distribution of adhesions along the nucleus-tip axis. **D)** Lifetime of GFP-paxillin positive adhesions in migrating leader astrocytes (top). Duration time of assembly and disassembly of these adhesions (bottom). **E)** Lifetime of GFP-paxillin positive adhesions in migrating astrocytes (left) of the second and third rows. Duration time of assembly and disassembly of these adhesions (right). **F)** Immunofluorescence images of astrocytes in the migrating monolayer stained for nuclei (cyan) and paxillin (green). Data is from N = 3 independent experiments. The sample size for each repeat is: si ctl 49, 50, 50, si triple IF 50, 50, 50 and si plectin 50, 50, 55 for 3C; si ctl 26, 40, 40 and si triple IF 26, 40, 40 for 3D, si ctl 16, 16, 35 and si triple IF 8, 36, 37 for 3E. P values are n.s. (not significant) for p > 0.05, * for p < 0.05, **: p for p < 0.01 and *** for p < 0.001. Scale bar 5 μm for 3A and 10 μm for 3B and 3F.

We thus asked whether IFs could affect FAs as previously reported in vimentin-expressing cells (Burgstaller et al., 2010; Gregor et al., 2014). In migrating monolayer of control astrocytes, FAs are strongly concentrated at the leading edge of the leader cells (Fig. 3B, 3C), while only few small FAs are visible further back in the protrusion or at the rear of leader cells and in followers (Fig. 3B-D). Depletion of IF proteins increased the number and the size of FAs in leader cells (Fig. 3B, 3C, S3A). In this condition, FAs appeared more dispersed between the front edge and the nucleus (Fig. 3B (insets), 3C). Depletion of any single IF protein had a similar effect (Fig. S3B). Using GFP-paxillin expressing astrocytes, we observed that in leader cells as well as in follower cells of the second or third rows, the lifetime of FAs was longer in si triple IF cells than in control cells (Fig. 3D, 3E and movie 8). This reduced FA turnover resulted from both a slower assembly and a slower disassembly rate (Fig. 3D, 3E). Further away from the wound, the turnover of FAs was too slow to be analysed, however, we noticed that IF depletion also led to a strong increase of FAs (Fig. 3F), which is in agreement with the higher tractions seen in the centre of the monolayer of si triple IF cells (Fig 1F-I). Since microtubules, which are tightly associated with IFs, can affect the dynamics of FAs (Akhmanova et al., 2009; Ezratty et al., 2005; Gan et al., 2016), we analysed the impact of the si triple IF on microtubule organisation and dynamics. In si triple cells, the microtubule network was intact and globally oriented along the axis of migration like in control cells (Fig. S3C). We analysed microtubule dynamics by tracking the plus end protein, End-Binding protein 3 (EB3). We observed that IF depletion slightly but significantly modified the speed and the directionality of EB3 comets (Fig S3D and movie 9) indicating that IFs impact microtubule dynamic instability. This also confirms that loss of IFs has a slight effect on microtubule orientation, as previously reported (Gan et al., 2016). Thus, we conclude that the effect of IF depletion on FA dynamics does not result from major changes in the microtubule network but may involve alterations of microtubule dynamics and the control of microtubule associated proteins (Chang et al., 2008; Jiu et al., 2017).

In migrating astrocytes, the cytoskeletal crosslinker plectin colocalises with IFs and paxillin at the distal end of FAs (Fig. 4A). Although we could not distinguish between the different plectin isoforms, plectin1f is the most likely candidate as it has been shown to anchor vimentin to FAs and control FA turnover in fibroblasts (Burgstaller et al., 2010). IF interaction with plectin and with FAs was confirmed by co-immunoprecipitations. Vimentin and GFAP also co-immunoprecipitated with plectin, talin and vinculin but were not found in control immunoprecipitations performed with irrelevant IgGs (Fig. 4B). Comparison of immunoprecipitations performed on non-migrating and migrating cells (4h after wounding) showed that IF association with plectin and FA proteins strongly increased during wound-induced migration (Fig. 4B).

**FIGURE 4:**
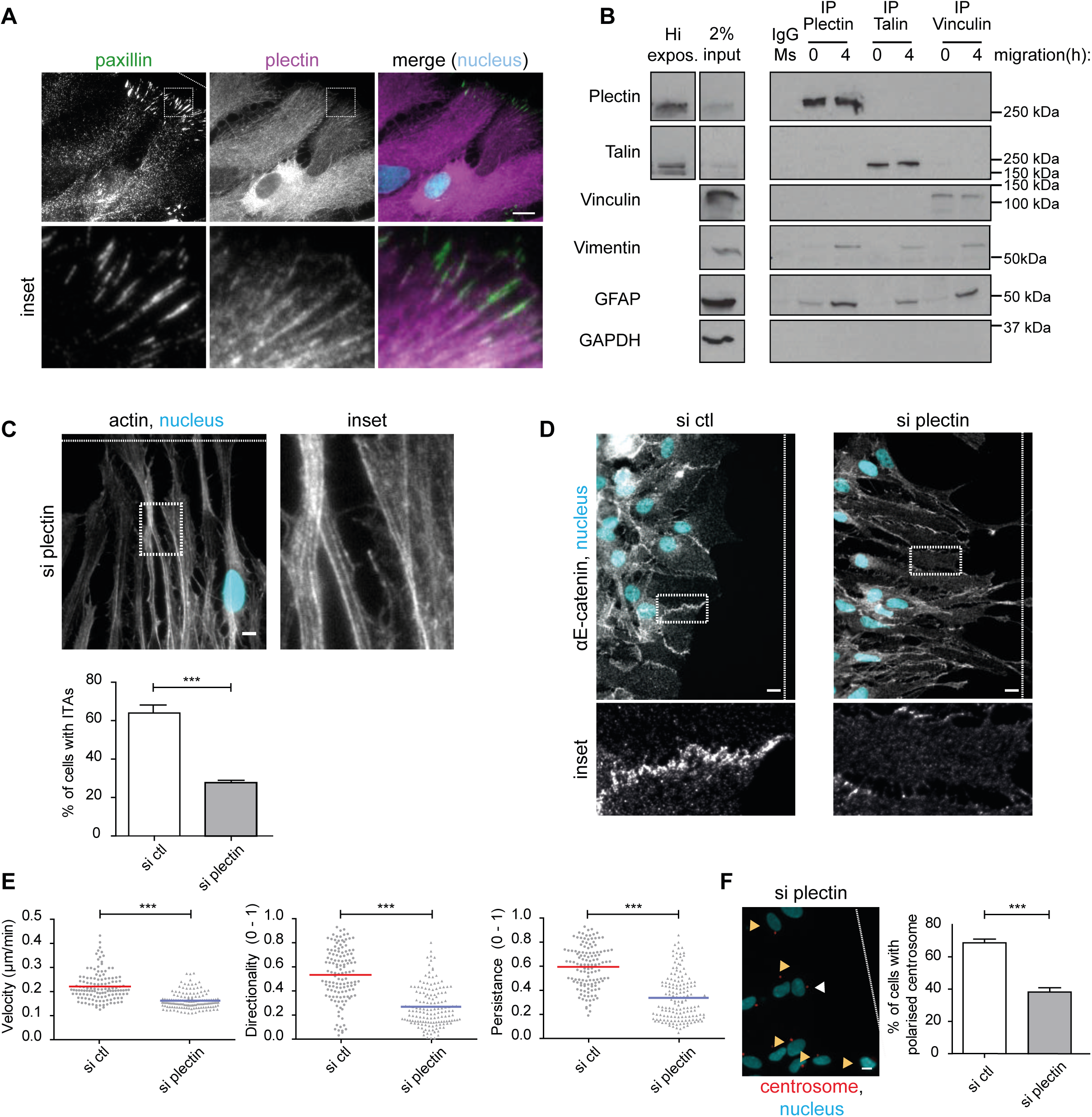
Plectin knockdown phenocopies IF depletion. **A)** Immunofluorescence images of migrating astrocytes stained for nuclei (cyan), paxillin (green) and plectin (magenta). **B)** Western blot analysis using indicated antibodies of total astrocyte lysates (left panels) and of proteins immunoprecipitated with antibodies against plectin, talin or vinculin and control antibodies (IgG Ms) before and 4h after wounding. Input lysate corresponds to 2% of the total lysate used for the IP, with a higher exposition shown for plectin and talin bands. **C)** Immunofluorescence images of migrating astrocytes stained for nuclei (cyan) and actin (phalloidin, grey) compared to si ctl (Fig. 2A). The white dotted line represents the position of the wound. The quantification shows the percentage of cells that present ITAs. **D)** Immunofluorescence images of migrating astrocytes stained for the AJ marker α-E-catenin (grey) and nuclei (cyan). The white dotted line represents the position of the wound. **E)** Quantification of nuclear tracking of migrating astrocytes after 24h of migration. The graphs show cell velocity, directionality and persistence of migration of astrocytes nucleofected with si ctl or si plectin. **F)** Immunofluorescence images of centrosome orientation in plectin-depleted astrocytes stained for nuclei (cyan) and centrin (red) compared to si ctl (Fig. S1F). White arrowheads were scored as polarised centrosomes, while yellow ones were scored as non-polarised centrosomes. The graphs show the percentage of cells with the centrosome located in the wound-facing quadrant in front of the nucleus. Data is from N = 3 independent experiments. The sample size for each repeat is: si ctl 262, 247, 150 and si plectin 384, 245, 155 for 4C; si ctl 40, 50, 39 and siplectin 50, 50, 49 for 4E; si ctl 92, 222, 83 and si plectin 101, 210, 123 for 4F. P values are *** for p < 0.001. Histograms show mean ± s.e.m. Scale bar 10 μm.

We then investigated the role of plectin in astrocyte migration using siRNA to reduce plectin expression in primary astrocytes (Fig. S3E, S3F). Plectin depletion strongly affected the distribution of FAs, that are not confined only to the cell tip (Fig. 3B, 3C). It also affected actin organisation (presence of stress fibres but not ITAs, Fig. 4C) and AJs (loss of junctions, Fig. 4D), mimicking the effects observed following IF protein depletion, confirming observations made on plectin-depleted endothelial cells (Osmanagic-Myers et al., 2015). Plectin depletion reduced the speed, directionality and persistence of migration (Fig. 4E) as well as centrosome orientation (Fig. 4F). further supporting the idea that plectin and IFs act together to control AJ and FA dynamics, actin organisation and collective cell migration (Leduc and Etienne-Manneville, 2015; Leube et al., 2015; Seltmann et al., 2015; Wiche et al., 2015)}.

### IFs control vinculin-mediated traction forces at FAs

Since IFs strongly influence actin organisation and the generation of traction forces in the migrating monolayer, we asked whether IF depletion may affect the localisation of vinculin. Vinculin is a key mechanosensor protein controlling the interaction of actin with both FAs and AJs (De Pascalis and Etienne-Manneville, 2017; Gregor et al., 2014). In most control cells, vinculin was found both at FAs and AJs (Fig. 5A, 5B). In contrast, in a large majority of IF-depleted cells, vinculin was found in FAs but was frequently absent from AJs (Fig. 5A, 5B). Vinculin exists in a closed conformation which opens upon stretching when actin fibres pull on the carboxy-terminal domain of talin-bound vinculin (Bakolitsa et al., 2004). We took advantage of the vinculin FRET tension sensor (Grashoff et al., 2010) to quantify the tension exerted on vinculin during cell migration. The FRET index distribution in FAs was shifted to statistically lower values in si triple IF leader cells than in control leader cells (Fig. 5C), suggesting that vinculin-mediated tension was increased in IF depleted cells.

**FIGURE 5:**
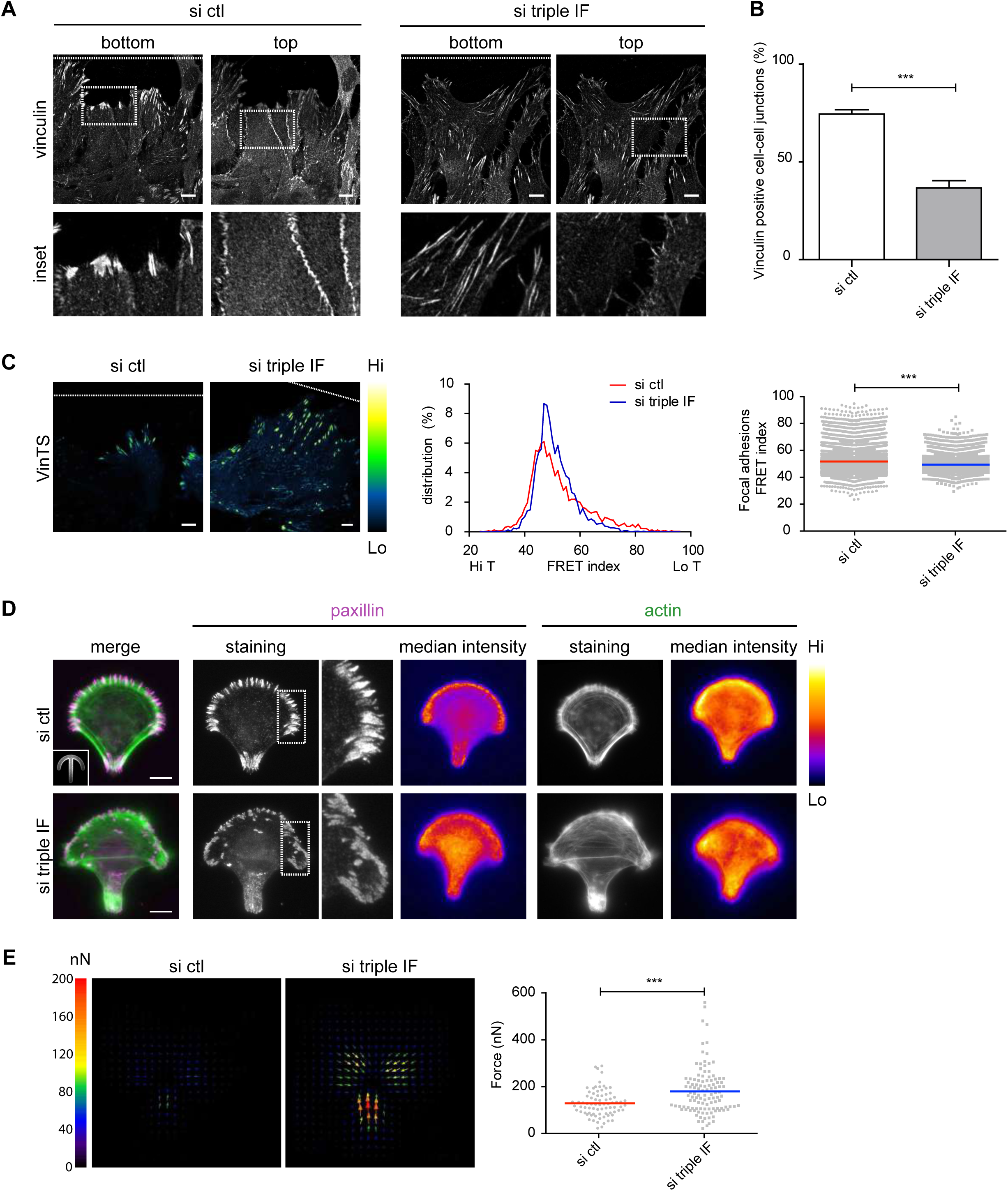
Intermediate filaments limit the generation of traction forces at focal adhesions. **A)** Immunofluorescence confocal images in two distinct focus planes of astrocytes stained for vinculin. The white dotted line indicates the position of the wound. **B)** Graph showing the percentage of junctions showing a positive vinculin staining. **C)** Fluorescence images of migrating astrocytes expressing VinTS (Green Fire Blue LUT). Representative frequency distribution of frequencies of FRET index (central graph) and quantification of the mean FRET index (inversely related to vinculin tension) of vinculin in focal adhesions (right graph). Each dot represents a focal adhesion. The white dotted line indicates the position of the wound. **D)** Immunofluorescence and intensity projection (fire LUT) of paxillin (magenta) and actin (green) on micropatterned cells. Inset shows the crossbow shaped micropattern. Projections are calculated on one experiment representative of three. **E)** Left, tractions fields of astrocytes plated on collagen coated crossbows. Arrows indicate the orientation, colour and length of the local magnitude of the force in nN. Right, quantification of traction forces in single micropatterned cells. Data is from N = 3 independent experiments. Histograms show mean ± s.e.m. The sample size for each repeat is: si ctl 137, 247, 150 and si triple IF 222, 200, 151 for 5B; si ctl 2537, 3312, 1473 adhesions and si triple IF 1447, 2548, 2081 adhesions for 5C; si ctl 32, 21, 25 and 30, 46, 38 for 5E. P values for *** is p < 0.001. Scale bar 10 μm.

Since AJs have been shown to locally inhibit FA formation (Camand et al., 2012), the effect of IFs on FAs may result from the alteration of AJs leading to a redistribution of FAs and actin. To test this hypothesis, we plated single cells on adhesive micropatterns, which allowed us to study the role of IFs on FAs and traction forces independently of cell migration and cell-cell interactions. On crossbow-shaped micropatterns, FAs reproducibly concentrated on the outer edge of the arc and on the rear side of the bar forming the pattern, while the actin cable systematically formed on the two non-adhesive edges of the cell (Fig. 5D). IF depletion led to globally wider distribution of FAs, which appeared on the inner region of the adhesive stripes while actin cables formed in more random direction across the cell body (Fig. 5D). We then used TFM to analyse the orientation and strength of traction forces of single cells plated on crossbow shaped micropatterns. IF depleted cells exerted significantly stronger traction forces than control cells (Fig. 5E). These results show that IFs impact FAs, actin organisation and traction forces independently of their effect on AJs and cell migration.

In conclusion, our results show that glial IFs increase migration speed, direction and persistence during astrocyte collective movement. Keratin-depletion has an opposite effect on keratinocyte migration (Osmanagic-Myers et al., 2006) strongly suggesting that IF functions on cell migration depend on their composition. Although the silencing of IF protein is only partial, it strongly affects collective migration showing that limited perturbations of the IF network as observed in IF-related diseases or oncogenesis are sufficient to alter cell functions. We show that glial IFs together with plectin control the distribution of FAs, the organisation of the actin cytoskeleton in the migrating cell sheet as well as on the single cell. The glial IF network controls the distribution of tractions forces generated in the migrating cell sheet, preventing the accumulation of such forces within the monolayer. It is thus likely that IF-depleted followers are less pulled by the leader cells, and instead essentially migrate in response to the tractions they exert on the substrate, leading to an alteration of the collective behaviour. This could partially be due to the fact that IF depleted glial cells cannot interact with their neighbours, as a similar effect was observed for keratin 8 morphants of *Xenopus* mesoendoderm explants (Sonavane et al., 2017). In agreement with this hypothesis, we show here that IFs promote the formation of ITAs, which connect the lateral sides of the adjacent leaders and control the treadmilling of AJs. Such actin-driven AJ dynamics is required for the maintenance of cell-cell contacts, which control the directionality and the persistence of migration (Camand et al., 2012; Peglion et al., 2014). However, experiments of single cells indicate that the control of AJs by IFs is not solely responsible for the impact of IFs on FAs and traction forces. The IF associated protein, plectin, interacts with IFs, localises at FAs and its depletion mimics most of the effects observed after IF depletion. IFs may act *via* plectin to control vinculin recruitment and the mechanical coupling between FAs and actin and thereby traction forces and/or act indirectly by controling the acto-myosin network (Costigliola et al., 2017) or regulating FA dynamics through microtubules (Akhmanova et al., 2009; Ezratty et al., 2005; Gan et al., 2016).

Altogether, these observations provide new insights into how higher IF levels may contribute to the invasive phenotype of glioblastoma, suggesting that IFs may serve as an interesting therapeutic target to reduce the invasive capacity of these tumours.

## Materials and Methods

### Cell culture

Primary astrocytes were obtained from E17 rat embryos (Etienne-Manneville, 2006). Use of these animals is in compliance with ethical regulations and has been approved from the Prefecture de Police and Direction départementale des services vétérinaires de Paris. Astrocytes were grown in 1g/L glucose DMEM supplemented with 10% FBS (Invitrogen, Carlsbad, CA), 1% penicillin-streptomycin (Gibco) and 1% Amphotericin B (Gibco) at 5% CO_2_ and 37° C.

### Transfection

Astrocytes were transfected with Lonza glial transfection solution and electroporated with a Nucleofector machine (Lonza). Cells were then plated on appropriate supports previously coated with poly-L-Ornithine (Sigma). Experiments are carried out 3 or 4 days post-transfection and comparable protein silencing was observed. siRNAs were used at 1 nmol and DNA was used at 5 *μ* g (or 2.5 *μ* g for pLifeAct-mCherry). siRNA sequences used were: Luciferase (control) UAAGGCUAUGAAGAGAUAC, GFAP rat GAGUGGUAUCGGUCCAAGU, GFAP rat set B CAACCUGGCUGUGUACAGA, Vimentin rat UGAAGAAGCUGCACGAUGA, Vimentin rat set B CGAGCUCAGCACAUAACAA, Nestin rat GUUCCAGCUGGCUGUGGAA, Nestin rat set B AUAAGAGCCUUCUAGAAGA, Plectin rat: AGAGCGAGCUGGAGCGACA. DNA plasmids used were: pEGFP-paxillin (gift from E. M. Vallés, Institut Curie, Paris), pPaxillin psmOrange (Addgene 31923), pPaxillin-mCherry cloned from pPaxillin psmOrange with BamHI and NotI (gift from M. Piel, IPGG, Paris), pEGFP-N3-Vimentin (Sakamoto et al., 2013), mCherry-N3-Vimentin cloned from pEGFP-N3- Vimentin, pVimentin psmOrange (Addgene 31922), pLifeAct-mCherry (gift from M. Piel, IPGG, Paris), pEGFP-N-cadherin (gift from C. Gauthier-Rouvière, CRBM, Montpellier) eGFP-RLC Myosin II (gift from B. Latge and B. Goud, Institut Curie, Paris), vinculin tension sensor (Vin TS, Addgene 26019 (Grashoff et al., 2010)) and mEB3-FL pEGFP-N1 (gift from N. Morin, CRBM, Montpellier).

### 2D wound healing assay

Cells were plated on appropriate supports (dishes, plates, coverslips or glass-bottom MatTek) and allowed to grow to confluence. Fresh medium was added the day after transfection and the day prior to the experiment. The cell monolayer was then scratched with a p200 pipette tip to induce migration.

### Immunofluorescence

Cells migrating for 8h (unless otherwise stated) were fixed with cold methanol for 3-5 min at -20° or 4% warm PFA for 10 min and permeabilised for 10 min with Triton 0.2%. Coverslips were blocked for 30 min with 4% BSA in PBS. The same solution was used for primary and secondary antibody incubation for 1h. Nuclei were stained with DAPI or Hoechst and were mounted with Mowiol. Epifluorescence images were acquired with a Leica DM6000 microscope equipped with 40X 1.25 NA or 63X 1.4 NA objectives and recorded on a CCD camera with Leica software. Confocal images were acquired with a Zeiss LSM 700 inverted microscope equipped with a 63X NA 1.3 oil objective. Super-resolution 3D-SIM images were acquired with a Zeiss LSM780 ELYRA with 63X 1.4 NA or 100X 1.46 NA objectives and recorded on an EMCCD camera Andor Ixon 887 1K with Zen software.

Primary antibodies used: anti- *α* -E-catenin Abcam 32635 (lot 4 of 04/2016), anti-*a*-tubulin Biorad MCA77G YL1/2, anti-GFAP Santa cruz sc-6170 (clone C-19, lots F0313 and H189) and Dako Z0334 (lot 20035993), anti-N-cadherin Abcam ab12221 (lot GR139340-26) and sc-31030 (clone K-20, lot B2014), anti-Nestin Millipore MAB353 (lot 2780475), anti-paxillin BD 610051 (lot 5246880) and Abcam ab32084 (clone Y133 and lot GR215998-1), anti-Phalloidin Abcam 176759 (lot GR278180-3), anti-pericentrin Covance PRB 432-C, anti-plectin Abcam Sigma (clone 7A8), anti-talin Sigma T3287 (clone 8D4, lot 035M4805V), anti-vimentin Santa cruz sc-7557R (clone C-20, lot E3113), 7557 (clone C-20, lots E0313 and B1510) and Sigma V6630 (lot 10M4831), anti-vinculin Sigma V9131 (lot 036M4797V). Secondary antibodies were all purchased from Jackson ImmunoResearch.

### Live imaging

For phase contrast wound healing assays, cells were wounded in 12 well plates. HEPES was added to the medium and paraffin was used to cover the well to prevent medium evaporation. Acquisition started 30 min after wounding. Movies were acquired with a Zeiss Axiovert 200M equipped with a thermostatic humid chamber with 5% CO_2_ and 37° C. All images were acquired with dry objective 10X 0.45 NA and an EMCCD camera. Images were acquired every 15 min for 24h. Nuclei of leader cells were manually tracked with Fiji software (Manual Tracking plugin). Calculations for 24h of migration were normalised by pixel size and time with the following formulas (*n* is the number of time points acquired, *θ* is velocity):

Average velocity

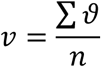

Persistence

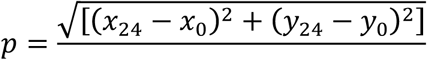

Directionality

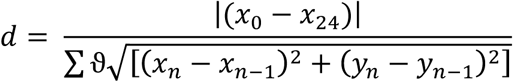

For fluorescence live imaging, cells on appropriate dishes (Ibidi or MatTek) were wounded and allowed to migrate 4h before image acquisition. HEPES and antioxidants were added to the medium before acquisition. Epifluorescence experiments investigating paxillin turnover, N-cadherin and actin retrograde flow were performed with a Nikon BioStation IM-Q (40X objective) at a rate of 1 frame/5 min for 3-5h with 5% CO_2_ and 37° C. Confocal experiments for MRLC and EB3 were performed on a Perkin-Elmer spinning disk confocal microscope equipped with an EMCCD camera and a 63X 1.4 NA objective with 5% CO_2_ and 37° C. Super-resolution experiments were acquired with the Zeiss LSM780 ELYRA at 37° C. Images were automatically processed for Structured Illumination with Zen software.

### FRET (Fluorescence Resonance Energy Transfer) tension sensor experiments

Cells were transfected with VinTS. The day prior to the experiment, medium was changed to phenol-free (FluoroBrite DMEM, Gibco A18967). Cells were wounded for 4h and HEPES was added before acquisition. Images were acquired with a Zeiss LSM780 with a 63X 1.4 NA objective and Definite Focus at 5% CO_2_ and 37°C. The donor was excited at 458 nm and the emission spectrum of the sensor was in 13 contiguous channels from 465 to 545nm, to include both the donor and acceptor emission wavelengths. Images were analysed through the Fiji plugin PixFret. After background correction, the FRET index was calculated on thresholded images to delineate the FAs. PixFRET calculates the FRET index by measuring the ratio of intensity I_FRET_/(I_Donor_+I_FRET_) between the donor channel and the FRET channel (channels 3 and 6).

### FACS (Fluorescence-activated cell sorting)

Migrating and non migrating cells in 100 mm dishes were trypsinised and fixed in 4% warm PFA for 10 min. 10,000 cells were analysed on a BD LSRFortessa II flow cytometer applying a live cell gate based on FSC (forward scatter) and SSC (side scatter).

### Micropatterns

Coverslips were plasma-cleaned for 45s and incubated immediately with 0.1 mg/ml PLL-PEG diluted in 10 mM HEPES for 30 min at room temperature. Excess of PLL-PEG was allowed to slide off and the coverslips were dried before printing. Micropatterns were printed for 4 min with specifically designed chrome masks and coated for 1h at 37° C with collagen (100 *μ* g/ml) and fluorescent fibrinogen (Alexa 647, 20 *μ* g/ml) diluted in NaHCO_3_ pH 8.3 100mM. Micropatterned coverslips were washed twice in PBS and used immediately for the experiments. Plated cells were allowed to adhere for 16h before fixation. Crossbow patterns have a diameter of 40 *μ* m, with an area of around 1000 *μ* m^2^.

### TFM (Traction Force Microscopy) experiments on micropatterns

After printing, micropatterns were coated with collagen and fluorescent fibrinogen for 1h. A thin layer of polyacrylamide solution (52.5 *μ* l acrylamide, 50.25 *μ* l bis-acrylamide, 381 *μ* l water, 10 μl 0.2 μm diameter fluorescent beads (FluoSpheres from Molecular Probes), 5 μl APS 10%, 3 μl Temed for a 40 kPa gel), previously vacuumed, was allowed to polymerise on the collagen for 20 min. Gels were then washed twice in NaHCO_3_ pH 8.3 100mM and cells are plated immediately. 16h after, stacks of single cells were acquired with a spinning disk microscope before and after trypsin treatment. Only single cells, checked through Hoechst staining, were analysed. Analysis of TFM on micropatterns was carried out with a custom-designed macro in Fiji based on the work by (Tseng et al., 2012). Briefly, the top-most planes of beads before and after trypsinisation were selected and aligned using a normalised cross-correlation algorithm (“Align slices in stack” Plugin). The displacement field was computed from bead movements using PIV. Parameters for the PIV were 3 interrogation windows of 128, 96 and 64 with a correlation of 0.60. Traction forces were calculated from the displacement field using the Fourier Transform Traction Cytometry (FTTC) and a Young Modulus of 40 kPa, a regularisation factor of 10-9 and a Poisson ratio of 0.5.

### TFM experiments on migrating cells

Transfected cells plated on 35 mm diameter plates were incubated with trypsin for 5 min and resuspended in 400 *μ* l of fresh media. They were plated in rectangular PDMS patterns on collagen-coated hydrogels of a Young Modulus of 9 kPa containing fluorescent beads (0.2 *μ* m diameter, FluoSpheres from Molecular Probes). Removal of the PDMS stamp allowed the cells to migrate outwards on the gel. Images were acquired every 15 min for 24 h on an inverted epifluorescence Nikon Eclipse Ti microscope with an ORCA-Flash 4.0 CMOS IMCD camera, 10X dry objective NA 0.3 with PFS at 5% CO_2_ and 37° C. Analysis was carried out every 1h for the 24h time course with custom-made Matlab software. For a more detailed protocol, see (Serra-Picamal et al., 2015; Tambe et al., 2011; Trepat et al., 2009).

### Electrophoresis and Western blot

Cells lysates were obtained with Laemmli buffer composed of 60 mM Tris-HCl pH6.8, 10% glycerol, 2% SDS and 50 mM DTT with the addition of anti protease (cOmplete cocktail, Roche 11 873 588 001), 1 mM glycerol phosphate, 1 mM Na orthovandate and 1 mM Na fluoride. Samples were boiled 5 min at 95° before loading on polyacrylamide gels. Transfer occurred at 0.2 or 0.3A overnight on nitrocellulose membranes. Membranes were blotted with TBST (0.2% Tween) and 5% milk and incubated 1h with the primary antibody and 1h with HRP-conjugated secondary antibody. Bands were revealed with ECL chemoluminescent substrate (Pierce, Thermo Scientific).

Primary antibodies used: anti- *²3* -catenin BD 610154 (lot 3137536), anti-GFAP Santa cruz sc-6170 (clone C-19, lots F0313 and H189), anti-GAPDH Millipore MAB374 (lot 2689153), anti-Nestin Millipore MAB353 (lot 2780475), anti-plectin Abcam Sigma (clone 7A8), anti-talin Sigma T3286 (clone 8D4, lot 035M4805V), anti-vimentin Sigma V6630 (lot 10M4831), anti-vinculin Sigma V9131 (lot 036M4797V). Secondary HRP antibodies were all purchased from Jackson ImmunoResearch.

### Immunoprecipitation

Confluent 100 mm diameter dishes of WT astrocytes were washed with cold PBS and lysed with IP B buffer (25 mM Tris HCL pH 7.5, 1 mM EDTA pH 8, 150 mM NaCl, 1 mM MgCl2, 0.5% Triton X-100) with the addition of fresh anti-protease (cOmplete cocktail, Roche 11 873 588 001). Lysates were centrifuged at 13.000 rpm for 2.30 min at 4°C. 15 μl (2% of the total lysate) of supernatant were stored at -20°C with the addition of Laemmli buffer while the rest of the supernatant was incubated with 50 μl of Protein G (previously washed in PBS) and 1 μg of primary antibody or control antibody for 2h at 4°C on a spinning wheel. Beads were washed at least 8 times with the IP B buffer before loading in Laemmli buffer on precast gels (Invitrogen).

### Quantifications of immunofluorescent images

All images were analysed with Fiji software (Schindelin et al., 2012). For actin distribution and orientation, cells in the first row were scored manually as positive when containing both ITAs and SF (interjunctional transverse arcs and stress fibres) or negative when they possessed only SF and no ITAs. To measure actin fibre orientation in the followers, phalloidin z-stacks were projected for maximum intensity onto one image. After filtering with a specific kernel and despeckled, they were analysed with a specific Fiji directionality plugin. Vinculin localisation on focal adhesions or cell-cell junctions was manually scored in the first row of cells. Polarity was also scored by counting the percentage of cells with the centrosome (centrin staining) oriented in the 90° quadrant towards the wound.

To quantify retrograde flow, kymographs (width = 5 pixels) were drawn on the cell-cell junctions (front part) or on the central axis (tip and behind) of the cell for myosin. They were plotted with the Multi Kymograph plugin from which three slopes were measured to obtain an average value for each cell. To quantify focal adhesions, background subtracted images of focal adhesion stained cells were thresholded with the same value for all images. Cell contours were drawn manually to count the number and size of focal adhesions. The relative projected distance between focal adhesions and the nucleus was defined to characterise the distribution of focal adhesions by projecting the FA position onto a line drawn between the centroid of the nucleus and the tip of the cell (see schematics in Fig. 3C). To quantify paxillin focal adhesion turnover, integrated fluorescent intensity was analysed. Turnover was defined as the time elapsed between their birth (first frame) and their death (last frame). Assembly was defined as the time between the birth and the peak of fluorescent intensity, while disassembly was defined as the time between the highest peak of intensity and the death of the focal adhesion.

EB3 tracking was performed with Fiji TrackMate Plugin with the following settings: LoG detector, estimated blob diameter 1 pixel, simple LAP tracker (1 pixel linking max distance, 1.5 pixel gap-closing max distance and 3 gap-closing max frame gap) and filters for duration of track (more than 4) and track displacement (more than 2.5). Velocity, persistence and directionality towards the wound were calculated with the formulas previously described for cell migration (in the x direction) with a custom-designed Fiji macro.

### Statistical Analysis

All data is presented as the mean in dot plots or mean +/- s.e.m in histograms of at least three independent experiments. Statistical analysis was obtained with Student’s t-test for Fig. 2E, 2G, 3E, 3F, 4E, 5C, 5E and Supplementary Fig. 1A, 1C, 1D, 1E, 3D and 3E, ANOVA (ANalysis Of VAriance) followed by Bonferroni *post-hoc* test for Fig 1C, 2F, 3C, 3E, 3F and Supplementary Fig 2F or **x**^2^ test for contingency on the raw data for Fig. 2B, 2H, 4C, 4F, 5B, and Supplementary Fig 1F, 2C and 3D. Analysis was performed with GraphPad Prism 5.0. P values are reported as n.s. (not significant) for p > 0.05, * for p < 0.05, **: p for p < 0.01 and *** for p < 0.001.

All data and codes are available upon reasonable request.

## Author contributions

CDP designed, performed experiments and wrote the paper, CPG assisted in the set up TFM and analysed results, SS performed part of revision experiments, BB assisted with biochemistry and discussions, MB assisted with the set up TFM on micropatterns, BV assisted with the set up TFM on micropatterns and designed a custom macro for analysis of TFM and EB3, CL designed the macro for FA analysis and assisted with SRM images, NB assisted with FRET microscopy analysis, XT assisted with TFM and discussions

SEM supervised the project, analysed the results and wrote the paper.

## Acknowledgments

This work was supported by l’Association pour la Recherche contre le Cancer (ARC), La Ligue contre le cancer and the Institut Pasteur. CDP was a scholar in the Pasteur – Paris University (PPU) International PhD program and received a stipend from Fondation pour la Recherche Medicale (FRM), Institut Pasteur and a short term EMBO fellowship. CPG acknowledges support from Fundació “la Caixa”.

We would like to thank members of the SEM, XT and MT labs for support and discussion, as well as JB Manneville and C Alibert for stimulating discussion and the careful reading of the manuscript. We gratefully thank N Festuccia for support and help with FACS. We gratefully acknowledge JY Tinevez and A Salles and the Imagopole of Institut Pasteur (Paris, France) as well as the France-BioImaging infrastructure network supported by the French National Research Agency (ANR-10 – INSB -04; Investments for the Future) and the Région Ile-de-France (program Domaine d’Intérêt Majeur-Malinf) for the use of the Elyra microscope. The authors greatly acknowledge the Nikon Imaging Centre, Institut Curie (Paris, France), member of the French National Research Infrastructure France-BioImaging (ANR10-INBS-04) and the ImagoSeine core facility of the Institut Jacques Monod, member of IBiSA and France-BioImaging (ANR-10-INBS-04) infrastructures.

## Movies

### Movie 1: Astrocyte migration is altered upon IF depletion

Phase contrast movie showing the wound-induced migration of monolayers of primary astrocytes transfected with si ctl and si triple IF. Time lapse movie (one frame every 15min for 24h) was acquired with a Zeiss Axiovert 200M microscope with a 10x objective. Scale bar 50 μm

### Movie 2-3: IFs affect tractions during astrocyte migration

eMigration of primary astrocytes transfected with si ctl and si triple IF **(2)** and their tractions T_x_ **(3)** during migration on a collagen-coated 9kPa hydrogel. Time lapse movie (one frame every hour for 24h) was acquired with a Zeiss Axiovert 200M microscope with a 10x objective. Scale bar 50 μm.

### Movie 4: IFs affect myosin retrograde flow

GFP-MRLC retrograde flow during si ctl and si triple IF astrocyte migration. Time lapse movie (one frame every 2min for 15min) was acquired with a spinning-disk microscope and a 63× oil objective. Arrow indicated the direction of migration. Scale bar 10 μm.

### Movie 5: IFs affect N-cadherin retrograde flow

N-cadherin retrograde flow during si ctl and si triple IF astrocyte migration. Time lapse movie (one frame every 5min for 2h) was acquired with a Biostation with a 40× air objective and Phase and EGFP channels. Scale bar 10 μm.

### Movie 6, 7: IF dynamics near focal adhesions

**Movie 6**: migrating astrocyte transfected with GFP-vimentin and paxillin-Orange shown in Figure 3A. Time lapse movie (one frame every 30s for 180s) was acquired with a super-resolution Zeiss LSM780 ELYRA with 63x oil objective and automatically processed for Structured Illumination. Scale bar 20 μm.

### Movie 7: migrating astrocyte transfected with GFP-paxillin and vimentin-cherry

Time lapse movie (one frame every 30s for 180s) was acquired with a super-resolution Zeiss LSM780 ELYRA with 63x oil objective and automatically processed for Structured Illumination. Scale bar 20 μm.

### Movie 8: IFs affect focal adhesion turnover

GFP-paxillin turnover in si ctl and si triple IF astrocyte migration. Time lapse movie (one frame every 5min for 3h) was acquired with a Biostation with a 40× air objective and Phase and EGFP channels. Scale bar 10 μm.

### Movie 9: IFs affect EB3 dynamics

EB3-GFP dyanmics in si ctl and si triple IF astrocyte migration. Time lapse movie (one frame every 5sec for 300sec) was acquired with a spinning-disk microscope and a 63× oil objective. Arrow indicated the direction of migration. Scale bar 10 μm.

## Supplementary materials

**Figure Supplementary 1.**
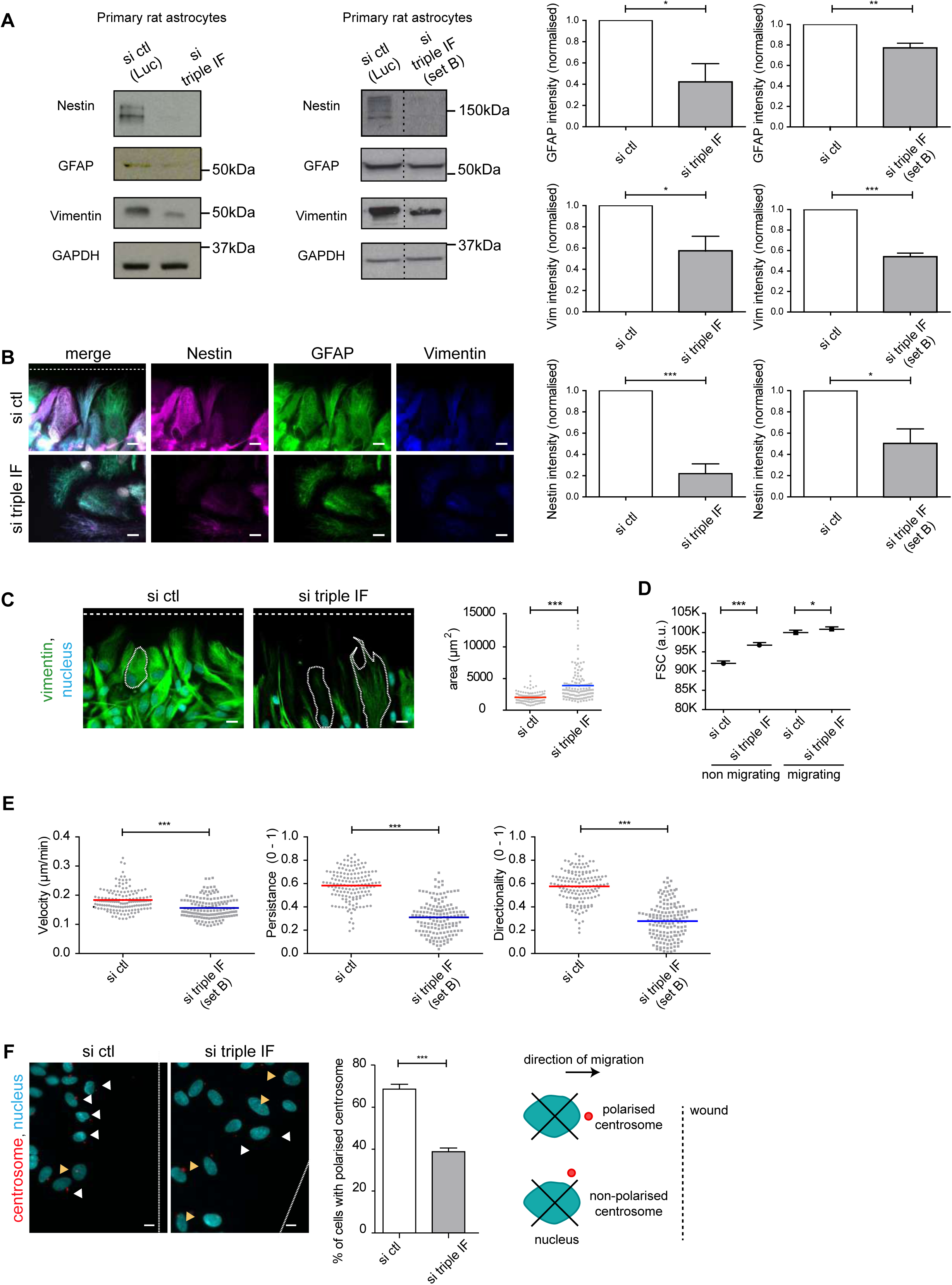
**A)** Western blots showing the expression levels of IF proteins in astrocytes (two sets) after 3 days of nucleofection with indicated siRNA. Graphs on the right show quantifications from three independent blots, normalised to si ctl and loading control. **B)** Immunofluorescence images showing Nestin (mouse Ab, magenta), GFAP (rabbit Ab, green) and vimentin (goat Ab, blue) in migrating astrocytes. The white dotted line represents the wound. **C)** Immunofluorescence of astrocytes stained for vimentin (green) and nuclei (cyan). The white dotted line represents the wound and examples of cell outlines. Quantifications show the spread area of cells during migration. **D)** Graph of FSC (Forward Scatter) proportional to the cell size, of migrating and non migrating cells in FACS analysis. **E)** Quantification of cell velocity, persistence and directionality after 24h of migration of astrocytes silenced for a second set of IF proteins (si triple IF set B). **F)** Immunofluorescence images of centrosome orientation in migrating astrocytes stained for nuclei (cyan) and centrin (red). The white dotted line represents the wound. White arrowheads represent polarised centrosomes, while yellow ones are non polarised. Scale bar 10 μm. The histogram shows the quantification of centrosome polarity (mean ± s.e.m.). Centrosomes are scored as polarised when they are found in the quadrant in front of the nucleus and facing the wound (right scheme). Random orientation of the centrosome with respect to the wound edge corresponds to a value of 25%. Cells analysed must be in the first row and must have neighbour cells on both sides. Data is from N = 3 independent experiments. The sample size for each repeat is: si ctl 30, 37, 50 and si triple IF 30, 37, 50 for S1C; si ctl 49, 49, 49 and si triple IF 50, 50, 50 for S1E; si ctl 137, 247, 150 and si triple IF 222, 200, 151 for S1E. P values are * for p < 0.05, **: p for p < 0.01 and *** for p < 0.001. Scale bar 10 μm.

**Figure S2.**
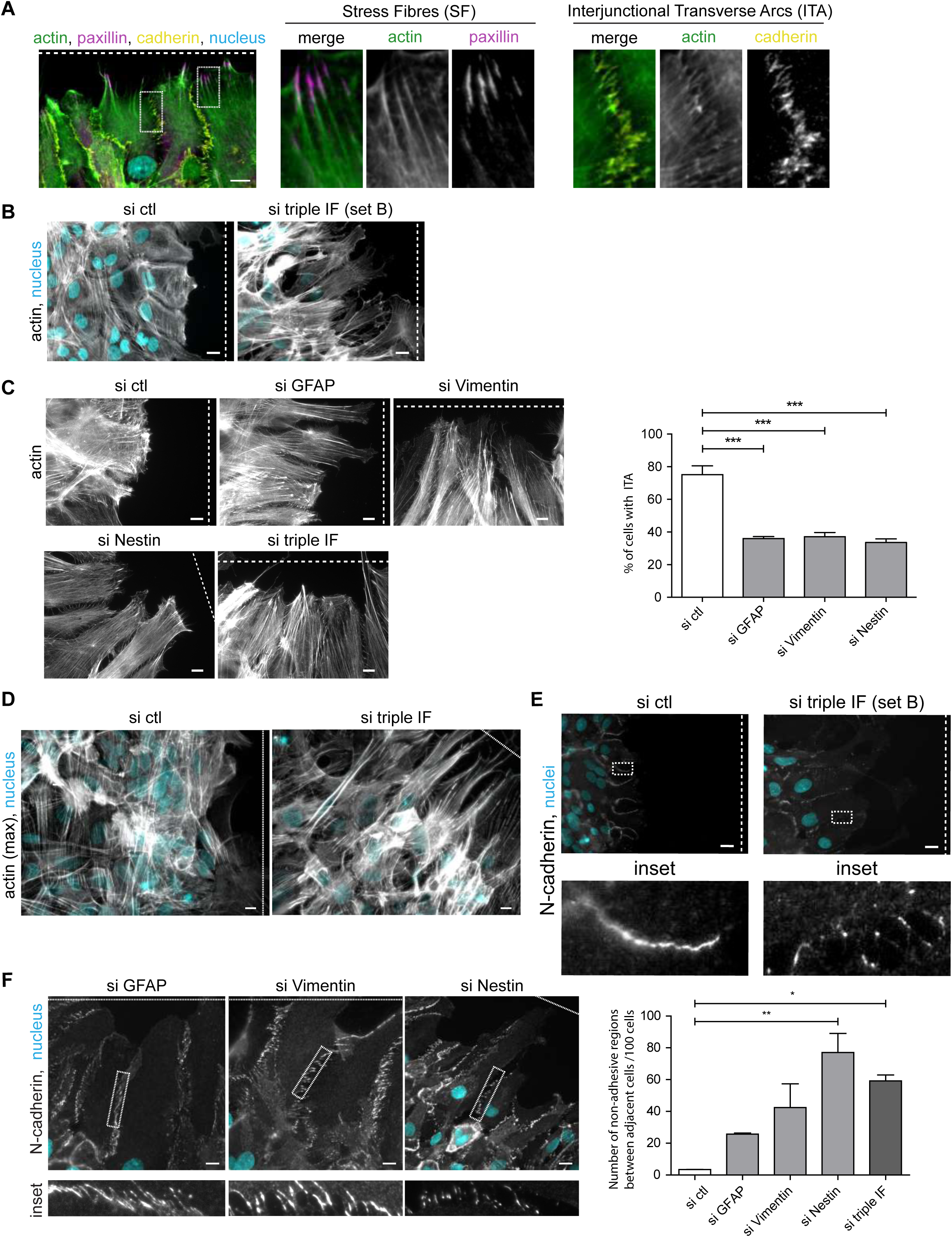
**A)** Immunofluorescence images of migrating astrocytes showing paxillin (magenta) for focal adhesions, cadherin (yellow, far red) for adherens junction, actin (phalloidin, green) and nuclei (cyan). **B)** Immunofluorescence images of migrating astrocytes nucleofected with the second set of si triple IFs (set B) and stained for nuclei (cyan) and F-actin (phalloidin, white). **C)** Immunofluorescence images of actin fibres in migrating astrocytes nucleofected with the indicated siRNAs and stained for F-actin (phalloidin, white). Graph shows the percentage of leader cells that present interjunctional transverse arcs (ITAs) in si control (si ctl) and si single depleted cells (si GFAP, Vimentin or Nestin). **D)** Maximal intensity projection of migrating astrocytes stained for nuclei (cyan) and actin (phalloidin, white) to show the orientation of actin in the rows behind the leading edge and used for the analysis of Figure 2C. **E)** Immunofluorescence images of migrating astrocytes nucleofected with the second set of si triple IFs (set B) and stained for nuclei (cyan) and N-cadherin (grey). **F)** Staining for N-cadherin (grey) and nuclei (cyan) in migrating astrocytes nucleofected with the indicated siRNAs compared to si ctl and si triple IF in figure 2H. In all images, white dotted line indicates the position of the wound. Data is from N = 3 independent experiments. The sample size for each repeat is: si ctl 71,145, 136, si GFAP 61, 131, 142, si vimentin 60, 135, 142 and si nestin 100, 131, 130 for S2C: si ctl 104, 120, 178 si GFAP 134, 137, 125, si vimentin 113, 180, 133, si nestin 132, 190, 123 and si triple IF 122, 225, 112 for S2F. P values are * for p < 0.05, **: p for p < 0.01 and *** for p < 0.001. Scale bar, 10 μm

**Figure S3.**
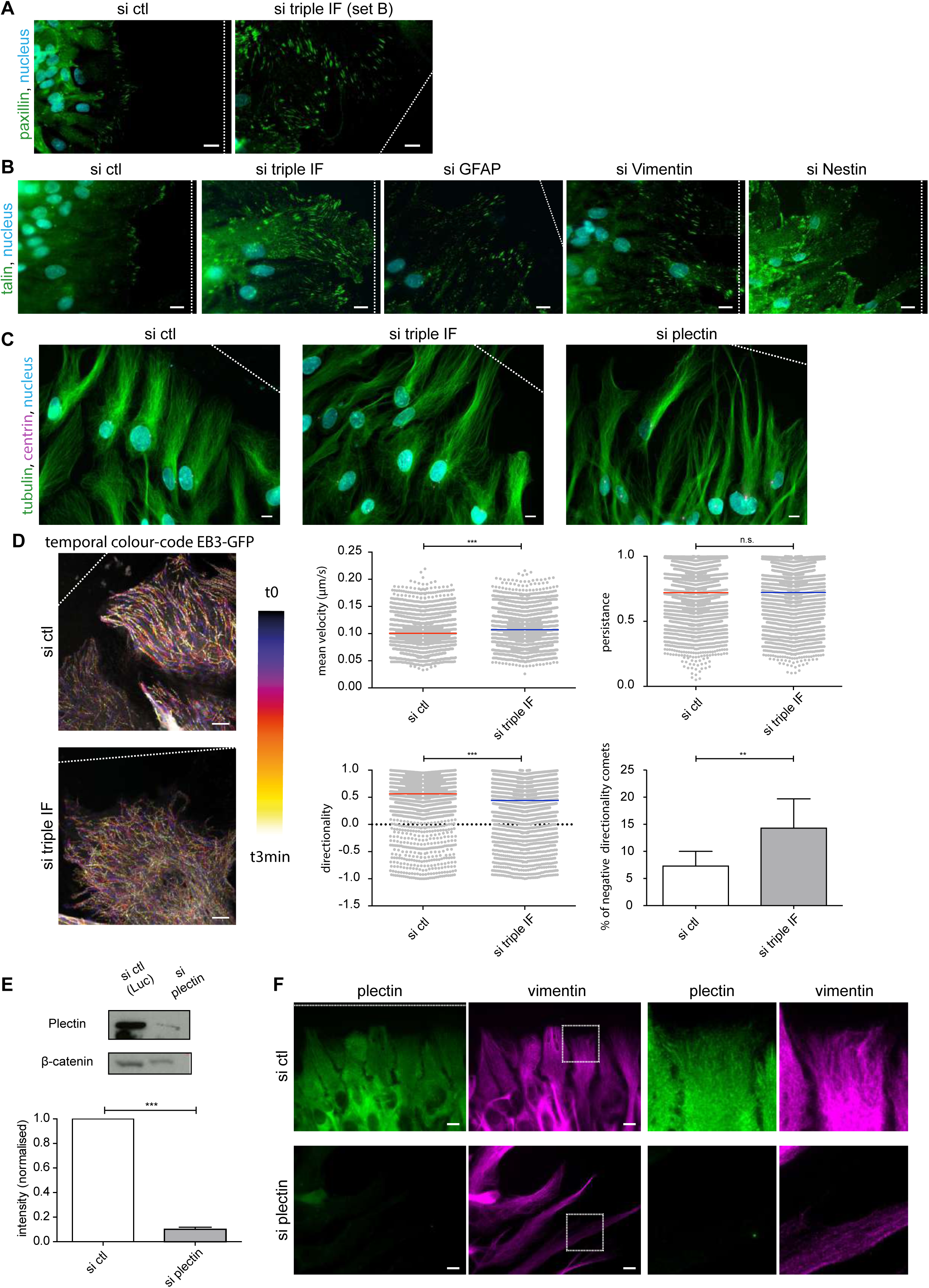
**A)** Immunofluorescence images of migrating astrocytes nucleofected with control siRNA (si ctl) or with a second set (set B) of si triple IFs and stained for paxillin (green) and nuclei (cyan). **B)** Immunofluorescence images of migrating astrocytes nucleofected with the indicated siRNAs and stained for talin (green) and nuclei (cyan). **C)** Immunofluorescence images of migrating astrocytes nucleofected with the indicated siRNAs and stained for tubulin (green), centrin (magenta) and nuclei (cyan). **D)** Temporal colour-code projection of EB3-GFP movies acquired on a confocal spinning disk (one image/5 s). Plots show mean velocity, persistence and directionality of EB3 comets compared to the wound. Histogram (mean ± s.e.m) shows the percentage of EB3 comets that have a positive directionality towards the wound. **E)** Western blot showing the expression level of plectin upon nucleofection with si plectin compared to si ctl. Histogram (mean ± s.e.m) shows the quantifications from three independent blots, normalised to si ctl and loading control. **F)** Immunofluorescence of migrating astrocytes stained for plectin (green) and vimentin (magenta). The sample size for each repeat is: si ctl 1650, 1108, 1378 and si triple IF 1652, 1035, 1331 (from at least 6 cells for experiment) for S3D. P values are n.s. (not significant) for p > 0.05, * for p < 0.05 and *** for p < 0.001. Scale bar 10 μm

## Abbreviations

AJ: Adherens Junction
EB3: End-Binding protein 3
FA: Focal Adhesion
GFAP: Glial Fibrillary Acidic Protein
IF: Intermediate Filaments
ITA: Interjunctional Transverse Arcs
PDMS: polymethylsiloxane
TFM: Traction Force Microscopy

